# Dielectrophoresis Reveals Stimulus-Induced Remodeling of Insulin Granule Subpopulations

**DOI:** 10.1101/2025.10.27.684914

**Authors:** Ashley Archambeau, Teji Korma, Aneesh Deshmukh, Mark A. Hayes, Kate L. White

## Abstract

The pancreatic β-cell contains several functional subpopulations of insulin secretory granules (ISGs). These subpopulations vary in maturity, age, and secretory capacity. Differences in protein and lipid composition of ISGs are correlated with disease but require further study to understand how ISG remodeling regulates normal biology. Due to limitations in traditional separation methods, the extent of these subpopulations, any overlap between them, and how they are affected by insulinotropic signals have not been determined. In this work, we adapted direct current insulator-based dielectrophoresis (DC-iDEP) to separate ISGs isolated from INS-1E cells, an immortalized rat insulinoma cell line model, according to their electrokinetic mobility ratio (EKMr). We were able to separate ISG subpopulations from unstimulated cells to determine a baseline distribution and identify characteristic profiles for immature, young, and old ISGs. We then analyzed distributions of subpopulations from cells stimulated with insulin secretion signals known to induce biophysical remodeling and maturation. We found significant changes in each subpopulation studied in response to stimulation, consistent with the increases in maturation, crystallization, and changes in size reported in the literature. This work provides new insights into how the cell controls ISG remodeling and may drive future development of more effective therapies.

**SIGNIFICANCE:** Understanding insulin secretory granule (ISG) heterogeneity and the functional role of subpopulations is a critical step towards unraveling the mechanisms of insulin secretion. We adapted direct current insulator-based dielectrophoresis (DC-iDEP) to resolve immature, young, and old ISG subpopulations based on their biophysical properties. We also explored the biophysical remodeling of ISGs under insulinotropic stimuli to further probe how the cell controls remodeling and secretion. Our work provides a new framework for quantifying granule heterogeneity by linking biophysical features to functional subtypes and assessing how environmental stimuli remodel ISGs. This methodology establishes a broadly applicable platform to interrogate organelle and vesicle diversity in complex biological systems.

## INTRODUCTION

Insulin is a peptide hormone essential for blood glucose regulation (1). It is produced by the pancreatic β-cell and packaged into organelles called insulin secretory granules (ISGs), which then undergo a process called maturation in which they are prepared for secretion upon β-cell stimulation (2). ISG maturation involves the acidification of the ISG lumen, conversion of proinsulin to insulin, and insulin condensation or crystallization (3,4). Due to the nature of this process, several ISG subpopulations with varying ages, cargo, secretory capacities, and levels of maturity are present in the β-cell, but the extent of the heterogeneity of ISGs in the β-cell is unclear (5-7). Disease states of the β-cell, such as diabetes, are associated with defects in maturation and changes in ISG subpopulation ratios (6,8). As rates of diabetes rise worldwide, understanding insulin maturation and ISG heterogeneity is crucial (9).

Several subpopulations have been identified. These subpopulations have been defined along multiple, overlapping axes, including functional differences, structural properties, and spatial arrangements within the cell. Functionally, ISG subpopulations have been described based on their secretion competence, calcium sensitivity, maturation state, and age (5,6,10-14). Structurally, ISGs exhibit heterogeneity in size, protein and lipid composition, and pH, properties that are tightly linked to functional behavior and maturation state (5,6,15-18). Spatially, ISGs vary in their proximity to the plasma membrane (PM) or mitochondria, including subpopulations such as the docked, readily releasable, or reserve pools, which are defined primarily by their distance from the PM (15,19-22). These functional, structural, and spatial categorizations are not independent. For example, granule age correlates with mobility, distance from the PM, and release competence (**Figure 1**), while composition and maturity levels influence calcium sensitivity and pH (5,6,11,18,23). For this reason, ISG subpopulations are often defined by overlapping features, complicating efforts to obtain a complete picture of biologically distinct subpopulations.

**Figure 1.**
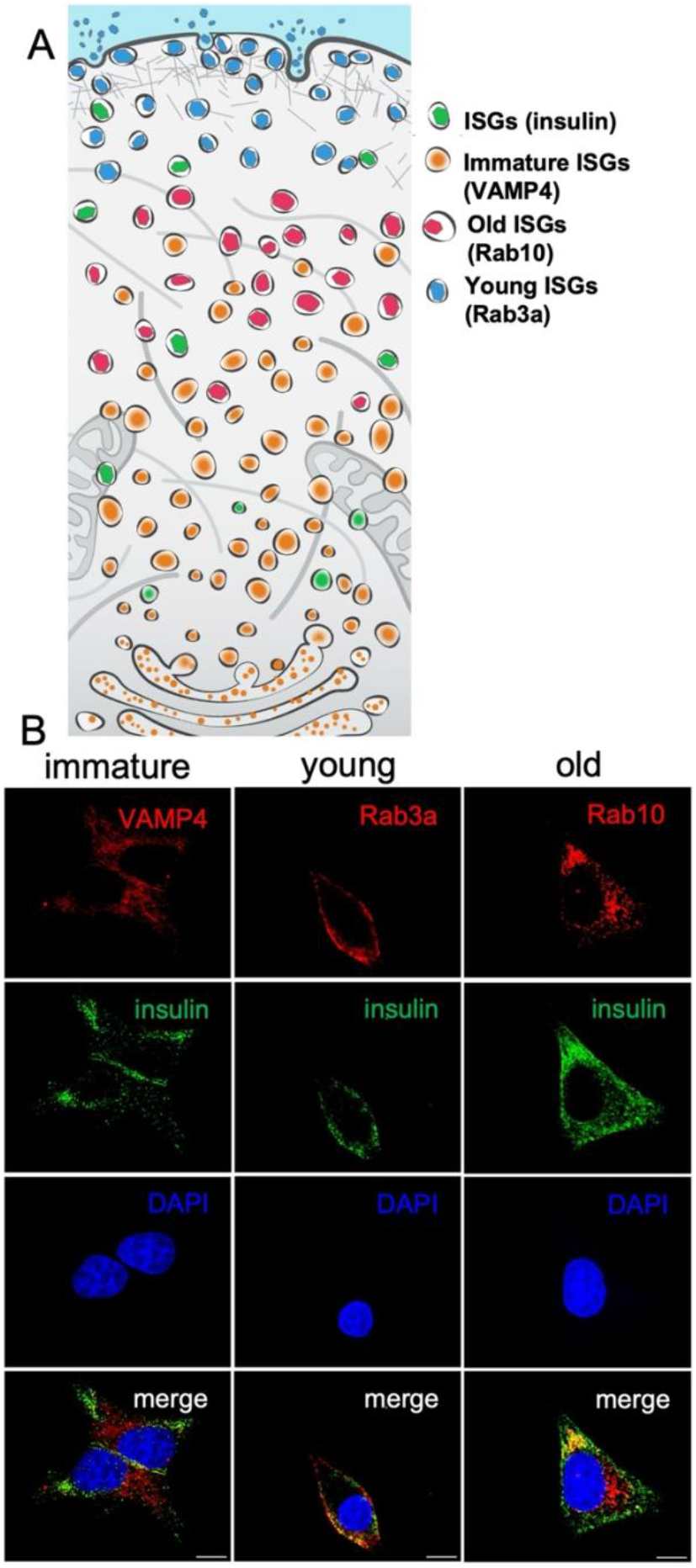
Insulin secretory granules form subpopulations.**A.**These subpopulations have characteristic spatial distributions and functions, with immature ISGs found near the interior of the cell, young ISGs primarily found near the PM, and older ISGs found at the interior of the cell while still colocalizing with insulin. **B**. Colocalization of ISG subpopulation markers VAMP4, Rab3a, and Rab10 with insulin in INS-1E cells. Scale bar: 5 µm.

Identification of subpopulations has traditionally required *a priori* knowledge of protein markers to distinguish subtypes using fluorescence microscopy. Alternative approaches of isolating ISGs for follow-up lipidomics and proteomic analysis provide highly heterogeneous results, likely due to differences in specific cellular conditions (7). It has been difficult to isolate ISGs for extensive biophysical characterization, as isolated ISGs often contain contamination by proteins from many other cellular compartments, such as the endoplasmic reticulum, trans-Golgi network, cytoskeletal, lysosomal, and mitochondrial proteins (7,24-31). To better understand the scope of ISG heterogeneity in the β-cell, an unbiased separation method is necessary. Dielectrophoresis has been used in the separation of both biological and nonbiological samples, such as proteins, stem cells, bacterial strains, and gold nanoparticles, among others (32-35). Thus, in our previous work, we established direct current insulator-based dielectrophoresis (DC-iDEP) as an avenue for unbiased separation of ISG subpopulations (36).

DC-iDEP separates particles using a nonuniform electric field based on their electrokinetic mobility ratio (EKMr). Biophysical differences between ISG subpopulations, particularly differences in radius, zeta potential or surface charge, and conductivity, affect the EKMr values of each subpopulation according to the equation below:

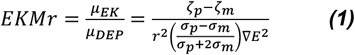

where *µ*_*EK*_ and *µ*_*DEP*_ refer to the electrokinetic and dielectrophoretic mobilities, *ξ*_*p*_ refers to the zeta potential of the particle, *ξ*_*m*_ refers to the zeta potential of the medium, *r* refers to the radius of the particle, *σ*_*p*_ refers to the conductivity of the particle, *σ*_*m*_ refers to the conductivity of the medium, and *E* refers to the electric field (37). Variations in the size, protein, and lipid compositions between different ISG subpopulations are expected to influence their EKMr distributions. A sawtooth-patterned microfluidic chip is used to accumulate particles, in this case ISGs, with specific EKMr values between the triangular tips, called “gates,” that have been designed to have a particular EKMr value at specific applied voltages (**Figure S1**) (32-35). Barekatain and Liu et al. used DC-iDEP to separate ISG subpopulations and reported significant changes in the ratios of ISG subpopulations between unstimulated and glucose-stimulated cells (36).

To build on this concept, we focused on ISG subpopulations defined by maturation stage, using known molecular markers, enabling us to assess the distributions of these subpopulations and their potential overlap. Immature, young, and old ISGs have been found to contain VAMP4, Rab3a, and Rab10, respectively (5,7,38) (**Figure 1A**). Accordingly, Rab3a, Rab10, and VAMP4-positive ISGs are hereafter referred to as young, old, and immature ISGs, respectively, and insulin-positive ISGs will be collectively referred to as ISGs. Immature ISGs are precursors to mature ISGs and are not typically secreted. Young, mature ISGs are more mobile, more acidic, and are preferentially released over older ISGs, while older ISGs are “caged” and targeted for degradation (5,11,18,39-42).

We also incorporate additional insulinotropic stimuli known to remodel ISGs and differentially affect ISG subpopulations, such as G-protein-coupled receptor 40 (GPR40) agonists and incretins (15). GPR40, a free fatty acid receptor, is involved in lipid signaling and the regulation of insulin secretion. Incretins are hormones that increase glucose-stimulated insulin secretion, ISG maturation, and include GLP-1 (glucagon-like peptide-1) and its analogs, and GIP (glucose-dependent insulinotropic polypeptide) (16,43). These two classes of drugs likely enhance ISG maturation differently, producing unique subpopulations (15,16). GPR40 agonists have been found to promote ISG maturation and to increase ISG diameter as the ISG approaches the periphery of the cell (16). The incretin exendin-4 has been found to promote ISG acidification and increase ISG density throughout the cell (16,18).

In this work, we measure the distribution and overlap of ISG subpopulations associated with differences in maturity and age in the INS-1E cell line. We compare the differences in this distribution as a result of orthogonal stimuli: a GPR40 agonist (TAK-875) and an incretin (exendin-4; Ex-4). We show that there are significant differences in the distributions of immature, young, and old ISGs, and that these distributions change in characteristic ways as a result of incretin stimulation and GPR40 agonism. Understanding how unique protein signaling pathways impact ISG maturation and generation of subpopulations is a critical step for unraveling the fundamental mechanisms of secretory biology and future development of effective therapeutics.

## EXPERIMENTAL METHODS

### Cell culture

INS-1E cells (a gift from Pierre Maechler’s laboratory at the University of Geneva) were cultured in 5% CO_2_ at 37°C. Cells were seeded in optimized RPMI 1640 media (AddexBio C0004-02, supplemented with 5% fetal bovine serum (FBS), 50 µM 2-mercaptoethanol, 1x Penicillin-Streptomycin, and 100 µg/mL streptomycin), sterile filtered through a 0.22 µm filter, and grown to 80% confluency. For each biological replicate, cells were seeded at a density of 4×10^4^ cells/cm^2^ in one-layer cell chambers (Avantor 734-1038). For stimulation, cells were starved in KRBH buffer containing 0 mM glucose for 30 minutes before stimulation in KRBH containing 25 mM glucose and either 10 nM exendin-4 or 10 µM TAK-875 for 30 minutes.

### Colocalization of subpopulation markers with insulin

Cells were grown on ibidi 8-well high ibiTreat slides (80806-96). Cells were fixed with 4% PFA for 10 minutes at 4°C, then stained with a primary antibody cocktail containing guinea pig anti-insulin (1:200, BioSynth 70R-10659) and an ISG subpopulation marker (mouse anti-Rab3a, 1:100 [Synaptic Systems 107 111]; mouse anti-insulin, 1:200 [Cell Signaling Technology 8138S]; rabbit anti-Rab10, 1:100 [Cell Signaling Technology 8127S]; rabbit anti-VAMP4, 1:100 [Synaptic Systems 136 002]) in 0.5% bovine serum albumin (BSA), 0.2% saponin, and 1% FBS in tris-buffered saline (TBS) at 4°C overnight. Excess antibody was removed by three washes with TBST (0.1% Tween-supplemented tris-buffered saline) for 10 minutes, then cells were stained with a secondary antibody cocktail containing Alexa Fluor® 488 AffiniPure™ Goat Anti-Guinea Pig IgG (H+L) (Jackson ImmunoResearch 106-545-003) and either Alexa Fluor® 568 AffiniPure™ Donkey Anti-Mouse IgG (H+L) (Jackson ImmunoResearch 715-575-150) or Alexa Fluor® 568 AffiniPure™ Goat Anti-Rabbit IgG (H+L) (Jackson ImmunoResearch 111-575-144) for 30 minutes at room temperature. Excess antibody was removed by 3 washes for 10 minutes, then cells were mounted in ProLong Glass Antifade Mountant with NucBlue Stain (Thermo Fisher Scientific P36981) and cured for 24 hours before imaging.

Confocal imaging was performed using a Leica Mica microhub equipped with a 63×/1.2NA water immersion objective. The signal was collected with 359, 499, and 650 nm excitations and 461, 520, 668 nm emissions for nuclei, insulin, and ISG subpopulation markers, respectively, by HyD FS detector. The images were deconvoluted by LAS X, lightning module.15-20 planes were collected with a Z-step of 0.16 µm, and a maximum projection of the planes was produced.

### Insulin secretory granule enrichment

Cells were harvested with 0.125% trypsin and gently washed in phosphate-buffered saline before being suspended in homogenization buffer (HB, 250 mM sucrose, 150 mM NaCl, 4 mM HEPES, pH 7.4, 1 mM EGTA) supplemented with house-made protease inhibitor (PI) cocktail (0.5 M AEBSF, 1 mM E-64, 1.13 mM leupeptin, and 151.36 μM aprotinin). Following trypsinization, all steps were performed at 4°C. Cells were homogenized by 20 strokes in a Dounce homogenizer (**Figure 2A**), then centrifuged at 1,500×g for 10 minutes to collect cell debris. Supernatant was collected, while the pellet was resuspended in HB supplemented with PI cocktail and subjected to homogenization and centrifugation as described above. The supernatants from both spins were then centrifuged at 5,900×g for 15 minutes to remove larger organelles. The supernatants from that spin were then pooled and centrifuged at 35,000×g for 60 minutes to sediment ISGs. The pellet was then resuspended in 400 µL HB and layered onto a density gradient column formed by two layers of Optiprep supplemented with 1 µM EDTA (Sigma-Aldrich D1556, 3.8 mL of 35% and 0.5 mL of 14.5%) in an open-top thin-wall polypropylene tube (Beckman 326819). The density column was centrifuged in an SW55i Beckman rotor at 190,000×g overnight to fractionate vesicle populations. The column-containing tube was punctured at the bottom, and 0.4 mL fractions were collected. Enzyme-linked Immunosorbent Assay (ELISA) was used to identify ISG-containing fractions. Similar dilutions were used for each fraction, and the manufacturer’s manual was followed for ELISA (Mercodia 10-1250-01). ISG-containing fractions were then pooled, diluted with HB, and centrifuged at 40,000×g for 60 minutes. The pelleted ISGs were resuspended in a low conductivity buffer (LCHB, 0.3 M sucrose, 5 mM MES, pH 6.3), which is compatible with dielectrophoresis studies. Enriched ISG samples were then confirmed to have the expected diameter by nanoparticle tracking analysis (NTA).

**Figure 2.**
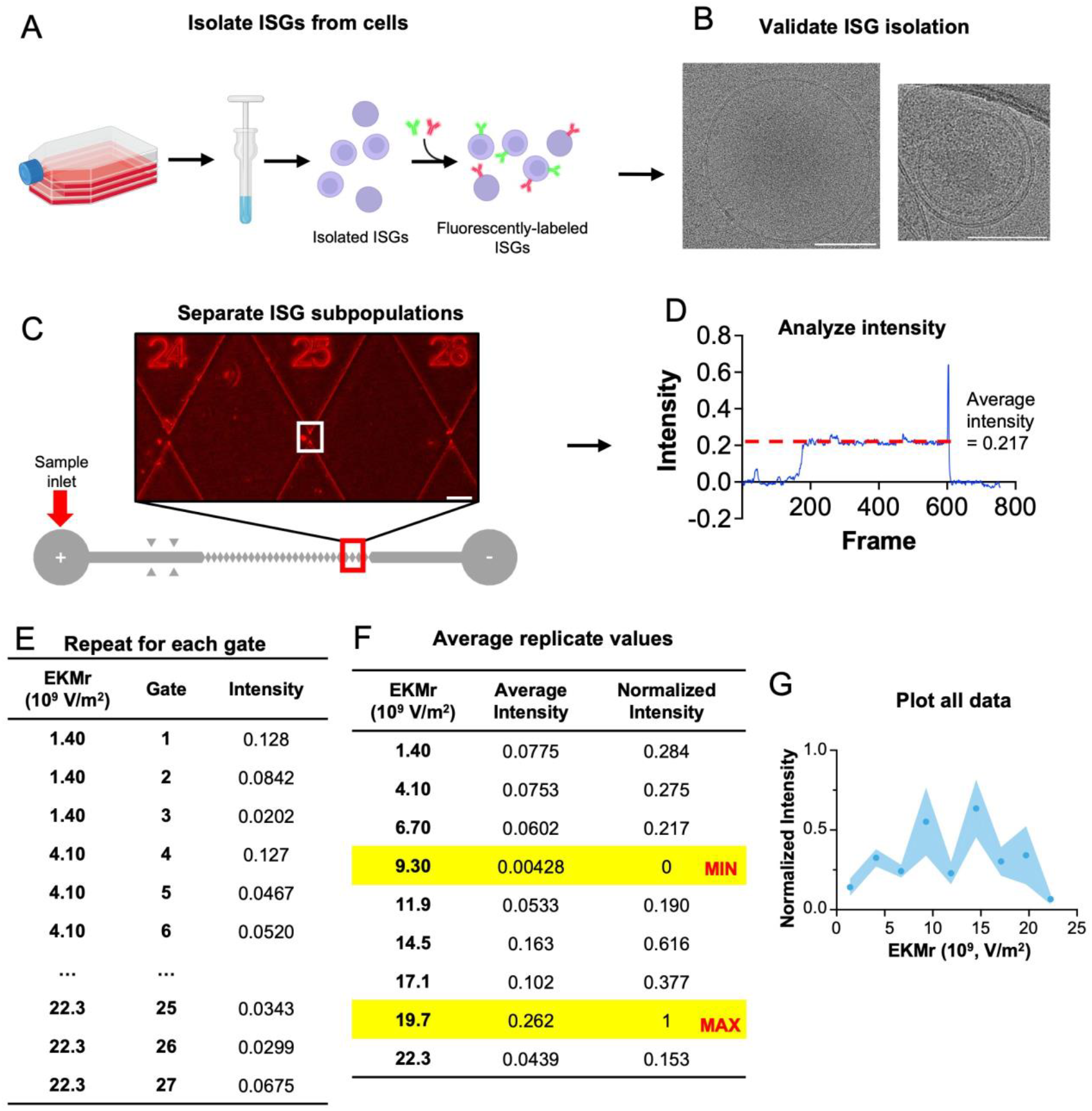
Workflow for separation experiments and analysis.**A.**ISGs were isolated from INS-1E cells using density gradient centrifugation, then pooling insulin-rich fractions (Created in BioRender). **B**. ISG isolation was validated using cryo-electron microscopy and other biochemical and biophysical analyses (pictured in **Figure S2**). **C**. Isolated ISGs were added to a sawtooth-patterned microfluidic channel and separated by applying a voltage. **D-F:** Normalization workflow for an example experiment. **D**. Intensity at the gate (white box in **E**.) is measured and normalized relative to the nearby channel and PDMS. **E**. The intensity of capture events is measured for each gate, with each technical replicate containing intensity measurements for each EKMr value in triplicate. **F**. The intensity measurements for each EKMr value are averaged and assigned values of 0-1 according to the ratio of the intensity to the difference between the maximum and minimum intensity values. **G**. Each replicate is plotted to provide the typical distribution of a subpopulation. Shown here is the distribution of Rab3a at 1500 V. Scale bars: **B**. 100 nm, **C**. 100 µm.

### Western Blotting

To determine the purity of the enriched ISGs, the cell pellet (post-1,500×g-centrifugation), the organelle pellet (post-5900×g-centrifugation), and the final pellet containing enriched ISG samples were mixed with 10× NuPAGE™ Sample Reducing Agent (Invitrogen NP0009) and 2× Novex™ Tris-Glycine SDS Sample Buffer (Invitrogen LC2676). Samples were loaded on Novex™ Tris-Glycine Mini Protein Gels, 10-20% (Invitrogen XP10205BOX) and run in a mini gel tank at 160 V for 60 minutes. Protein on the gel was then transferred to nitrocellulose membranes using min iBlot™ 2 Transfer Stacks (Invitrogen IB23002) in an iBlot 2 dry blotting device (Invitrogen IB21001). The membrane was blocked in 5% BSA in TBST, then incubated with antibodies against marker proteins for ER (SEC61B, Invitrogen PA3-015), endosomes (EEA1, Cell Signaling Technology 2411S), exosomes (CD63, Novus Biologicals NB100-77913), mitochondria (Cytochrome C, Novus Biologicals NB100-56503), lysosomes (LAMP2A, Cell Signaling Technology), and ISGs (synaptotagmin 9, Synaptic Systems 105 053) at room temperature for 5 hrs. Membranes were then washed three times with TBST, then incubated with Alkaline Phosphatase AffiniPure Goat Anti-Mouse IgG (H+L) (Jackson ImmunoResearch 115-055-003) and Alkaline Phosphatase AffiniPure® Alpaca Anti-Rabbit IgG (H+L) (Jackson ImmunoResearch 611-055-215) at room temperature for 1 hr. Membranes were then washed three times with PBST, and bands were visualized with 1-Step™ NBT/BCIP Substrate Solution (Thermo Fisher Scientific 34042).

### Visualization of ISGs by cryo-electron microscopy

All animal studies were conducted using procedures approved and conducted per Institutional Animal Care and Use Committee (IACUC) guidelines at the University of Southern California (Animal Use Protocol #21120). Mice were 2-3 months of age for experiments. Islets from LSL-Salsa6f mice (RRID: IMSR_JAX:031968) and human islets (RRID:SCR _014387, received from the Integrated Islet Distribution Program) were isolated and dissociated into single cells (44). Islet cells or INS-1E cells were deposited onto 200 mesh gold lacey carbon grids (Ted Pella 01894G) and allowed to adhere for 24 hrs in media. For stimulation conditions, cells were starved in KRBH buffer containing 0 mM glucose for 30 minutes before stimulation in KRBH containing 25 mM glucose and either 10 nM exendin-4 or 10 µM TAK-875 for 30 minutes. ISGs were isolated from INS-1E cells as described above. Excess buffer was removed from grids for 2 or 5 s for isolated ISGs or cells, respectively, at 37°C and 97% humidity, then grids were plunge frozen in liquid ethane using a Vitrobot Mark IV (Thermo Fisher Scientific). Grids were clipped using NanoSoft Autogrid rings and clips (MiTeGen M-CEM-NS-11011001).

Autogrids were screened under cryogenic conditions using a 200 kV Glacios Cryo TEM equipped with a Falcon 4 detector (Thermo Fisher Scientific, magnification 13,500× or 93,000×). Mouse and INS-1E cells were additionally imaged as part of tilt series on a 300 kV Krios G3i equipped with a Gatan K3 direct detection camera (Thermo Fisher Scientific, 26,000× magnification), but only frames at 0° tilt were analyzed in this work.

### Immunolabeling of ISGs

Isolated ISGs were incubated with 1:100 each of either anti-insulin (Cell Signaling Technology #8138) and anti-VAMP4 (Synaptic Systems 136 002) or anti-Rab3a (Synaptic Systems 107 111) and anti-Rab10 (Cell Signaling #8127) overnight. ISGs were washed with LCHB, then fluorescently labeled by incubation with 1:100 each of an Alexa 488-conjugated secondary antibody (Jackson ImmunoResearch 115-545-146) and an Alexa 568-conjugated secondary antibody (Jackson ImmunoResearch 111-575-144) for two hours. Fluorescently labeled ISGs were then washed once with LCHB to remove excess antibody and finally resuspended in LCHB.

### Device fabrication

The microfluidic device was fabricated and designed as described in previous publications (36,45). There are 27 gates (paired triangle tips) ranging in size from 25-73 µm, with wider gates closer to the inlet. The width of the gates decreases approximately 5 µm after every three repeats. Direct current was applied to the inlet and outlet, with potentials of 900 V, 1200 V, and 1500 V tested.

### Separation of ISG subpopulations by DC-iDEP

The separation channel was pretreated with 4% BSA for 15 minutes before washing with LCHB. 10-15 µL of an ISG sample was added to the inlet, and the volume in the channel was maintained at the outlet by the addition of LCHB to prevent pressure-driven flow. Direct current was applied at 900 V, 1200 V, and 1500 V using a 3000D High Voltage Sequencer (LabSmith HVS448), and particles were allowed to move through the channel according to their EKMr values (**Figure 2C**).

Separation experiments were imaged on a Leica DM IL LED Fluo Inverted Laboratory Microscope equipped with an S80/0.30 condenser and a 5×, NA 0.12 objective. Videos were recorded using a Leica Flexacam C1 camera at 1080p resolution and a frame rate of 30 frames per second. A Leica EL6000 external light source was used with a 3-position fluorescence slider for fluorescence excitation.

### Normalization

Intensity values were measured for the duration of each recording in three locations per gate: the gate, the PDMS near the gate, and an empty location in the channel using Fiji (46). Each list of intensity values was averaged during a capture event (particles present in the gate, **Figure 2D**). Gates containing no capture events were given intensity values of 0. The gate intensity when empty was subtracted from the intensity during a capture event, then normalized to the PDMS and channel (**Figure 2E**).

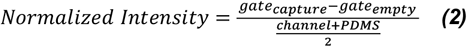

Gates with replicate EKMr values were first averaged. The resulting values were then normalized to a 0-1 scale based on the minimum and maximum intensities within each dataset to account for differences in concentration and fluorophore intensity (**Figure 2F)**. These final normalized values were plotted against their respective EKMr values (**Figure 2G**).

## RESULTS

### Localization of ISG subpopulations in INS-1E cells

We chose to focus on ISG subpopulations that reflect key stages of maturation and fluorescently labeled these subpopulations based on previously reported membrane markers. Upon fluorescently labeling ISG subpopulation markers for immature, young, and old ISGs, these subpopulations were found to localize to distinct regions of the cell with the expected localization of their respective subpopulation (**Error! Reference source not found.B**), as found in prior studies (5,11,18,47,48). VAMP4 (immature ISGs) did not colocalize with insulin due to proinsulin not being processed to insulin yet, and was primarily found far from the plasma membrane. Rab3a (young ISGs) was present largely at the periphery of the cell and had high colocalization with insulin, suggesting these ISGs are mature and ready for secretion (13,19,22). Rab10 (old ISGs) was mostly present in the interior of the cell, consistent with findings of old ISGs being caged in the interior of the cell (42,49).

### Analysis of ISGs post-isolation

Next, we isolated ISGs from INS-1E cells. Isolated fractions from density gradient centrifugation were analyzed by insulin ELISA. Fractions 8-10 were identified as ISG-containing fractions (**Figure S2A**) and were pooled and analyzed using a variety of biochemical and biophysical analyses (**Figure 2B; Figure S2**). Western blotting showed that ISG (synaptotagmin 9), lysosomal (LAMP2A), and exosomal (CD63) markers were enriched in pooled ISG samples. The presence of the exosomal marker CD63 in isolated ISGs is consistent with the finding of exosomes inside dense core vesicles but may also indicate the presence of contaminants (50). LAMP2A has also been shown to colocalize with ISGs, but its presence may be indicative of potential contamination (30). Fluorescently labeling ISG subpopulation markers prevented contaminating particles from being analyzed. ER, endosome, and mitochondrial markers were found in cell and organelle lysates but not enriched ISG samples (**Figure S2B**). NTA and CryoEM both confirmed the enriched ISG samples contained particles of the expected size (diameter: 200-500 nm, **Figure 2B; Figure S2C**) (19,51).

### Comparison of ISG subpopulation distributions in unstimulated cells

We then fluorescently labeled ISGs using ISG subpopulation-specific markers and separated them by DC-iDEP. Separation of ISG subpopulations by DC-iDEP revealed different EKMr distributions of ISG subpopulations (**Error! Reference source not found.A**). We used three applied voltages to separate ISG subpopulations by DC-iDEP: 900 V, 1200 V, and 1500 V (**Figure S3**). Each voltage affects the environment of the microfluidic channel differently, therefore changing the EKMr of each gate in the channel. The EKMr distributions are different at each voltage due to binning, as the microfluidic channel has gates with discrete rather than continuous EKMr values. With this channel design, lower voltages have greater separation at lower EKMr values, whereas higher voltages allow for better separation pf particles with higher EKMr values. Due to this binning, particles that are separate at lower voltages are captured at the same gate at higher voltages. Ultimately, we decided to focus on the 1500V datasets due to the broader EKMr range (**Figure S4**). Given the strong inverse relationship between EKMr and radius, particles with higher EKMr values are predicted to correspond to smaller ISGs.

Insulin was used as an overall ISG marker, and the superposition of each marker largely correlates with that of insulin (**Figure S5**), indicating that the three subpopulations encompass most ISG subpopulation states. The immature ISG marker VAMP4 exhibited a heterogeneous distribution, characterized by low-EKMr ISGs (7). Rab3a, a young ISG marker, had a bimodal EKMr distribution at mid-range values, while the old ISG marker Rab10 displayed ISGs at high EKMr values (**Figure 3A**) (5). Differences between subpopulation markers were analyzed via two-way ANOVA with Bonferroni post hoc correction (**Table S1**). Significant differences were found between young and old ISGs (9.30×10^9^ V/m^2^, p=0.0495; 1.45×10^10^ V/m^2^, p=0.0461; 2.23×10^10^ V/m^2^, p=0.0178), immature ISGs and insulin (4.10×10^9^ V/m^2^, p=0.0157), and immature and old ISGs (1.45×10^10^ V/m^2^, p=0.0489) (**Error! Reference source not found.B**). These differences indicate that each population has distinct distribution patterns due to the biophysical differences in ISGs, with overlap between distributions likely indicating overlapping subpopulations. This overlap between subpopulations is likely especially present within young ISGs, as this pool likely contains ISGs with a range of maturity levels.

**Figure 3.**
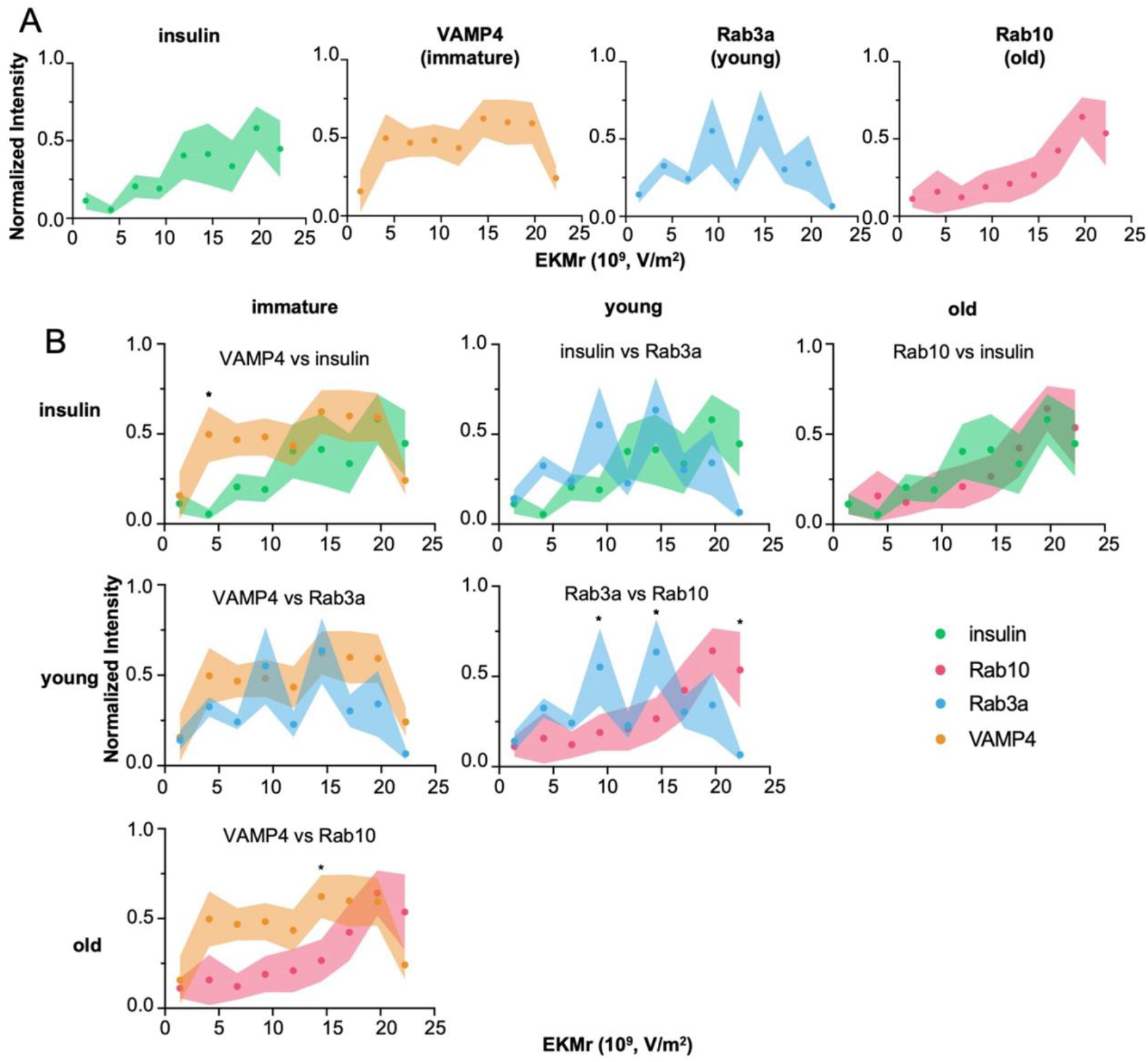
ISG subpopulations associated with different maturation stages have characteristic EKMr distributions. **A.**EKMr distributions of ISGs in unstimulated cells (n=3-4 biologically independent experiments). **B**. Comparisons of the distributions of different ISG subpopulations. Values are mean ± SEM (*p≤0.05 using ANOVA with Bonferroni post hoc multiple comparison correction).

### Effect of insulinotropic drugs on ISG subpopulations

To better understand how insulinotropic signals impact ISG remodeling or the shift in distinct subpopulations, we compared EKMr distributions of ISGs from unstimulated cells to those from cells stimulated by either TAK-875 or Ex-4. Both stimuli had significant effects on the EKMr distribution of isolated ISGs when analyzed via two-way ANOVA with Bonferroni post hoc correction, with TAK-875 producing larger shifts than Ex-4 across all applied voltages (**Figure S6**).

TAK-875 significantly decreased immature ISGs at several EKMr values, while Ex-4 did not produce significant changes at individual EKMr values. Specifically, immature ISGs were decreased by TAK-875 (4.10×10^9^ V/m^2^, p=0.0264; 6.70×10^9^ V/m^2^, p=0.0345; 1.45×10^10^ V/m^2^, p=0.0243; **Figure 4A**). Both TAK-875 and Ex-4 stimulation affected the shape of the EKMr distribution of immature ISGs. When stimulated by either TAK-875 or Ex-4, there was a smaller proportion of ISGs containing VAMP4 at lower EKMr values than in the unstimulated condition (**Figure 4A**). In contrast to the significantly decreased intensity of immature ISGs at 1.45×10^10^ V/m^2^ in response to TAK-875, Ex-4 stimulation led to a single peak in the EKMr distribution at the same EKMr value (**Figure 4A**), potentially revealing a condition-dependent maturation pathway producing mid-sized ISGs shared by both TAK-875 and Ex-4 stimulation.

**Figure 4.**
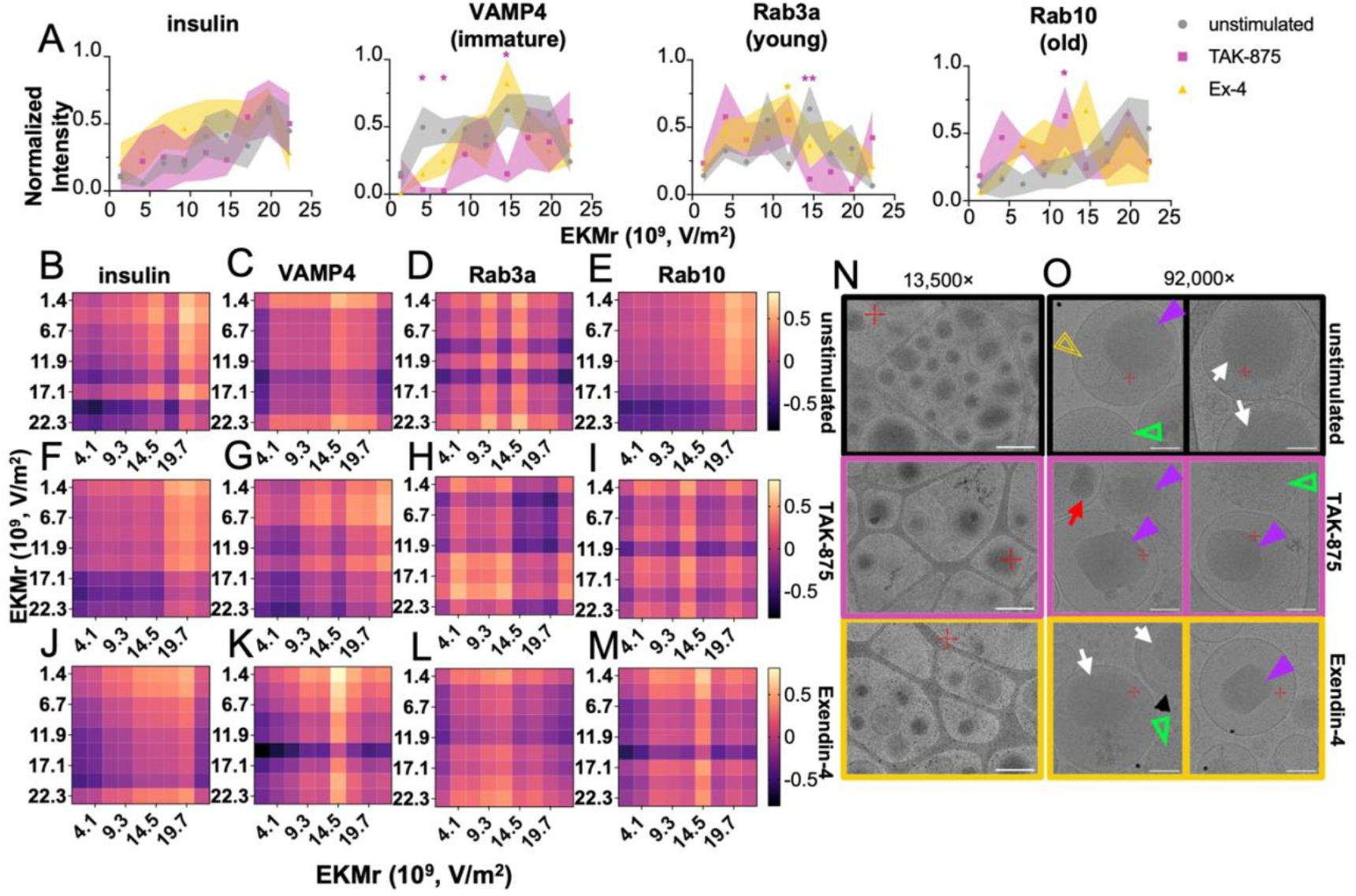
ISG subpopulations undergo remodeling in response to insulinotropic stimulation. **A.**EKMr distributions of ISGs isolated from unstimulated, TAK-875-stimulated (n=2 biologically independent experiments), and Ex-4-stimulated cells (n=2-3 biologically independent experiments). **B-M**. Heatmaps displaying the differences in average intensities between each pair of EKMr values within ISG subpopulation markers and **B-E**. unstimulated, **F-I**. TAK-875-stimulated, and **J-M**. Ex-4-stimulated conditions. **N-O**. Cryo-electron micrographs of ISGs inside unstimulated (top), Ex-4-stimulated (middle), and TAK-875-stimulated (bottom) mouse primary β-cells at **N**. 13,500x magnification, pixel size 2.06 nm, and **O**. 92,000x magnification, pixel size 0.15 nm. **O**. Symbols depict vesicles inside ISGs (yellow, double-lined arrowheads), crystalline cores (purple arrowheads), condensing cores (white arrows), immature ISGs (green, empty arrowheads), aggregates (red arrow), and fibrils (black arrow). Values are mean ± SEM (*p≤0.05, **p≤0.01 using ANOVA with Bonferroni post hoc multiple comparison correction. Comparisons made between stimulation and unstimulated condition, symbols color-coded according to condition). Scale bars: **N**: 500 nm; **O**: 100 nm.

Both TAK-875 and Ex-4 caused significant changes in the EKMr distribution of young ISGs at 1500 V (**Figure 4A**). TAK-875 induced a significant decrease in young ISGs at 1.45×10^10^ V/m^2^ (p=0.00960), while Ex-4 increased the abundance of ISGs at 1.19×10^10^ V/m^2^ (p=0.0441). TAK-875 and Ex-4 both eliminated the bimodal distribution of young ISGs observed in unstimulated cells, with TAK-875 favoring low EKMr values, consistent with a shift towards larger ISGs, and Ex-4 producing a more heterogeneous distribution (**Figure 4A**) (16).

Old ISGs also exhibited significant remodeling in response to stimulation. In the TAK-875 condition, old ISGs increased significantly at 1500 V (1.19×10^10^ V/m^2^, p=0.0417; **Figure 4A**), potentially corresponding to mid-sized ISGs. Notably, both TAK-875 and Ex-4 had significant effects on old ISGs at multiple EKMr values at both 900 V and 1500 V (**Table S2**; **Figure S6D**). Both TAK-875 and Ex-4 increased the proportion of old ISGs with lower EKMr values than in the unstimulated condition, with lower-EKMr peaks in the TAK-875 condition than in the Ex-4 condition.

To understand how multiple subpopulations may be affected similarly by the same condition that might not be obvious in the original plots, we visualized relative EKMr shifts using heatmaps generated from normalized intensity differences within a single subpopulation and condition (**Figure 4B-M**). These heatmaps allowed for the identification of other trends in EKMr distribution in response to stimulation. For example, in response to Ex-4, a similar peak at 1.45×10^10^ V/m^2^ in immature and old ISGs developed (**Figure 4A**), which is displayed in the heatmap by a dark horizontal or light vertical band (**Figure 4K,M**). Similarly, VAMP4 heatmaps show vertical bands at 1.45×10^10^ V/m^2^, indicating opposite effects on the relative amount of ISGs at the same EKMr value in response to TAK-875 and Ex-4 (**Figure 4G,K**). There were few significant differences between TAK-875 and Ex-4 stimulation, with Ex-4-stimulated cells having a significantly higher ratio of ISGs at 1.45×10^10^ V/m^2^ (p=0.00410), but the shapes of each were often different (**Figure S7, Table S3**).

As antibody binding could influence zeta potential and affect EKMr distributions, we examined whether the identity of a second antibody altered the EKMr distribution associated with young ISGs. ISGs labeled with Rab3a in combination with either VAMP4 or synaptotagmin 9 exhibited qualitatively similar EKMr distributions (**Figure S8**). No significant differences were detected between each dual-labeling scheme across EKMr values.

To visualize physical differences in ISGs under these different conditions, we imaged primary mouse β-cells using cryo-electron microscopy. There were observable differences in the appearance of ISGs in unstimulated cells compared with TAK-875 or Ex-4-stimulated cells. Unstimulated cells contained many ISGs with various levels of insulin condensation, as well as some ISGs containing exosome-like structures (50). As expected, based on soft X-ray tomography (SXT) experiments, the number of ISGs present near the cell periphery in TAK-875 and Ex-4 conditions was lower than in unstimulated cells (Error! Reference source not found.**N**) (16). TAK-875 stimulation caused higher levels of crystallization and more ordered crystals. In this condition, we also observed the presence of dark, unidentified aggregates attached to the cores of some ISGs that likely contributed to our previous observation of denser ISGs (Error! Reference source not found.**O**) (16). Exendin-4 stimulation led to increased crystallization, denser cores, and some ISGs with a smaller crystal-to-vesicle ratio than in the unstimulated condition (**Error! Reference source not found.N-O**).

## DISCUSSION

Subpopulations of ISGs have varying sizes and compositions, but the extent of these differences is unknown (5,6,15,16,18). In this work, our goal was to further develop separation methods and provide a more comprehensive analysis of the biophysical features of ISG subpopulations (36). In particular, we wanted to increase the interpretability of DC-iDEP separations by identifying the distribution of previously identified ISG subpopulations using Rab3a, Rab10, and VAMP4 antibodies to label young, old, and immature ISGs, as well as an insulin antibody to label all maturing ISGs (5,7). In this work, we found that these ISG subpopulations had characteristic distributions in unstimulated cells, indicating that known ISG pools exhibit distinct yet overlapping biophysical distributions (Error! Reference source not found.**A**).

### Interpreting the Size of ISGs and EKMr distributions

The traditional view in the field is that immature ISGs tend to be larger and less dense than mature ISGs. EKMr values vary inversely with radius (**Equation 1**) (16,52,53). This correlates with immature ISGs displaying higher ratios of ISGs at lower EKMr values, whereas young and old ISGs displayed distribution peaks at higher EKMr values. Insulin-positive ISGs exhibited two peaks at high EKMr values, likely reflecting enrichment of smaller, more mature ISGs (**Figure 3A**). These trends suggest that differences in ISG size contribute substantially to the observed EKMr distributions, although size alone cannot fully account for the observed heterogeneity.

### Zeta potential and conductivity of ISGs and EKMr distributions

In addition to size, differences in zeta potential and conductivity influence the EKMr value of a particle (**Equation 1**). Previous work has shown that young ISGs have lower zeta potentials and higher rigidities due to specific lipid compositions than older ISGs (5). These characteristics complement the distribution peaks at lower EKMr values of young ISGs than old ISGs in this work, with immature VAMP4 ISGs corresponding to low-EKMr values (**Error! Reference source not found.A**). Intravesicular pH may further contribute to EKMr differences, as mature ISGs exhibit lower luminal pH, which could increase conductivity and therefore contribute to shifts in EKMr distributions (18,54). Further biophysical characterization of the ISG subpopulations is necessary to completely understand which protein and lipid species are most influential in subpopulation formation.

### Heterogeneity within known subpopulations

In addition to the overall distribution trends, we observed substantial heterogeneity in the form of broad or multimodal EKMr distributions of ISGs within each subpopulation we studied. This indicates that there is appreciable heterogeneity in the biophysical makeup of these ISG subpopulations. Thus, each of the known subpopulations contains a collection of physically distinct ISGs. Insulin-labeled ISGs exhibited two major peaks that overlapped with the higher-EKMr peaks seen in young and old subpopulations, consistent with the insulin antibody labeling a broad range of maturing granules. We observed the most heterogeneity in immature and young ISGs, which may be due to activation of multiple maturation pathways giving rise to distinct subtypes over the lifetime of an ISG. VAMP4-positive ISGs had a broad distribution across the EKMr range tested, with a higher ratio of low-EKMr ISGs than other subpopulations (**Figure 3A, Figure S3C)**. The bimodal distribution of young ISGs potentially corresponds to the presence of young ISGs with a crystalline core (1.45×10^10^ V/m^2^) and young ISGs containing only condensed insulin (9.30×10^9^ V/m^2^), as the subset of ISGs positive for Rab3a likely contains ISGs in various stages of maturity (5). The heterogeneity observed within immature, young, and old subpopulations may be due to the presence of other previously described ISG pools, such as synaptotagmin 7 (syt7) or synaptotagmin 9 (syt9) subpopulations (**Figure S9**) (6). These subpopulations exhibit variations in their size, protein, and lipid compositions, which would affect their EKMr distribution, and are likely to exist in different maturation and age stages. In particular, the smaller radius of syt7 ISGs likely increases the average EKMr of these granules, but this effect is somewhat attenuated by the increased cholesterol increasing the rigidity of the membrane of syt7 granules (**Figure S9**) (6). Additional experiments probing these subpopulations and their changes in response to various conditions would provide valuable insight into ISG maturation.

### Impact of ISG maturation stimuli on ISG subpopulations

To better understand how different cellular pathways impact the generation of biophysically and functionally distinct ISG subtypes, we examined the effects of lipid signaling and incretin signaling on immature, young, and old ISGs. Two drugs that enhance ISG maturation through distinct signaling pathways, TAK-875 and exendin-4, have previously been reported to upregulate biophysically distinct ISG subpopulations (15,16,18). TAK-875 activates lipid signaling, and TAK-875 stimulation results in an increase in larger, denser ISGs that are often closer to the plasma membrane and distinct aggregate formations within the lumen (**Figure 4O)** (15,16). In contrast, Ex-4 stimulation causes more condensed crystalline cores throughout the cell (**Figure 4N-O**) (15,16).

Despite these differences, stimulation by TAK-875 or Ex-4 did not result in significant changes in the distribution of insulin-labeled ISGs, although Ex-4-treated cells exhibited a modest increase in the ratio of low-EKMr to high-EKMr insulin-labeled ISGs (**Figure 4A**). The sum of the EKMr distributions of these subpopulations does not overlay with the distribution of insulin as cleanly as in the unstimulated condition (**Figure S5**), suggesting that stimulation alters the relative abundance of ISG subpopulations rather than uniformly shifting all ISGs.

We saw a decrease in the subset of ISGs containing VAMP4 with low EKMr values in response to TAK-875 (**Figure 4A**), consistent with an increase in mature ISGs. Stimulation with TAK-875 also resulted in an increased proportion of low-EKMr young and old ISGs, accompanied by a reduction in immature ISGs compared to the unstimulated condition. This may indicate a rise in the radius of mature ISGs in response to TAK-875, consistent with previous studies. Stimulation with Ex-4 results in increased ISG maturation, decreased ISG diameter, and decreased ISG pH (15,16,18). We observed a decrease in low-EKMr VAMP4 ISGs, as well as a significant increase in young ISGs at 1.19×10^10^ V/m^2^, in response to Ex-4 (**Figure 4A**), indicating a decrease in immature ISGs and likely an increase in smaller, mature ISGs containing crystalline insulin.

Interestingly, both TAK-875 and Ex-4 stimulation were associated with modest but reproducible decreases in the abundance of old ISGs at 1.97×10^10^ V/m^2^. Although some changes in old ISG distributions could arise indirectly from the preferential secretion of young ISGs, β-cells are known to retain and cage older ISGs (11,18). Therefore, the observed shifts in old ISG distributions in response to TAK-875 and Ex-4 likely reflect some transformation or degradation of old ISGs rather than secretion alone. These decreases, while slight, may indicate remodeling of not only immature and newly formed ISGs, but also old ISGs.

The more pronounced effects of TAK-875 stimulation compared to Ex-4 stimulation may be due to lipid remodeling of the ISG membrane, which would be expected to alter the size and surface charges of ISGs (16). As the field learns more about inter-organelle contacts and communication, we will be able to piece together how these physical changes occur and specific lipids and proteins that are involved in regulating ISG maturation. Given that different cellular signals impact ISG biophysical characteristics, it is clear that there are multiple pathways for maturation and a diverse range of ISGs the cell can generate.

Future exploration of the effect of diseased states may improve our understanding of the functional implications of these changes, as certain subpopulations are known to be associated with either type 1 or type 2 diabetes (6). Identifying any overlap between the subpopulations described here and disease-related pools may enable a better understanding of these conditions and the development of future treatments. More broadly, this method may also prove useful in studying heterogeneity in other organelles or cells (55-61).

### Limitations of the study

A key limitation of separation by DC-iDEP is that ISGs must be removed from the cellular environment, which prevents direct correlation of EKMr values of an ISG with its native surroundings. Additionally, it can be challenging to isolate all ISGs for analysis. Immature ISGs lack the dense core that mature ISGs have and are found in different fractions than mature ISGs when isolated via traditional methods like density gradient columns (7). To address this challenge, future development of the separation channel to allow for the removal of separated subpopulations would enable the identification of more specific subpopulation markers. Labelling and imaging these specific markers *in cellulo* would assist in the interpretation of the identities of the separated subpopulations.

Additionally, modifying the channel to increase the resolution of mid-to-high EKMr values may improve the separation of ISG subpopulations. Because DC-iDEP provides higher sensitivity separations, further development as a preparative tool has powerful potential to methodically explore biophysical heterogeneity of ISGs, viruses, or other biologically complex particles.

INS-1E cells were used as a model of β-cells in this study for their ease of use and scalability, but the use of primary mouse or human β-cells may provide more insight into the heterogeneity of ISGs during disease progression. Primary β-cells appear to contain more crystalline insulin in their ISGs (**Figure S10**), which would likely affect EKMr distribution. They also contain other components that might affect the EKMr distribution, such as fibrils, aggregates, and vesicles (**Figure 4N-O**).

The markers we used likely label several types of ISG within the major subpopulations studied. Additionally, the antibodies themselves may modestly alter the EKMr of the ISGs they label. Because unlabeled ISGs cannot be analyzed in this system, we cannot directly quantify antibody-induced shifts. Comparisons using dual-labeling schemes suggest that relative differences between ISG subpopulations are robust to antibody choice (**Figure S8**). Each of these factors provides challenges in data interpretation, which may be made simpler by future technological advances. Despite these complications, these results provide new insights into the heterogeneity of ISGs and provide a basis for further use of DC-iDEP for exploring biological particles. Together, these findings establish DC-iDEP as a sensitive platform for resolving biophysical heterogeneity within organelle populations and provide a framework for linking physical properties to functional states.

## RESOURCE AVAILABILITY

### Lead Contact

**Kate L. White**

## AUTHOR CONTRIBUTIONS

Conceptualization, A.A. and K.L.W.; methodology, A.A., T.K., M.A.H., K.L.W.; investigation, A.A., T.K., A.D.; writing—initial draft, A.A. and K.L.W., writing—review & editing, A.A., K.L.W., T.K., A.D., M.A.H.; funding acquisition, K.L.W.; resources, K.L.W and M.A.H.; supervision, K.L.W. and M.A.H.

## ACKNOWLEDGMENTS

This material is based upon work supported by the National Science Foundation Graduate Research Fellowship Program under Grant No. DGE-1842487. Any opinions, findings, and conclusions or recommendations expressed in this material are those of the author(s) and do not necessarily reflect the views of the National Science Foundation. Funding from the National Institute of General Medical Sciences of the National Institutes of Health, under award number R35GM154893, and the USC Bridge Institute at USC provided financial support for this work. Thank you to Kevin Chang and Janielle Cuala for providing mouse islets for CryoEM experiments and Yekaterina Kadyshevskaya for illustration assistance. Thank you to Alex Ramirez for assisting in setting up the DEP device at USC. Human pancreatic islets were provided by the NIDDK-funded Integrated Islet Distribution Program (IIDP) (RRID:SCR_014387) at City of Hope, NIH Grant # U24DK098085. Images presented in this article were acquired at the Core Center of Excellence in Nano Imaging at the University of Southern California.

