## Supplemental Information for "Dielectrophoresis Reveals Stimulus-Induced Remodeling of Insulin Granule Subpopulations"

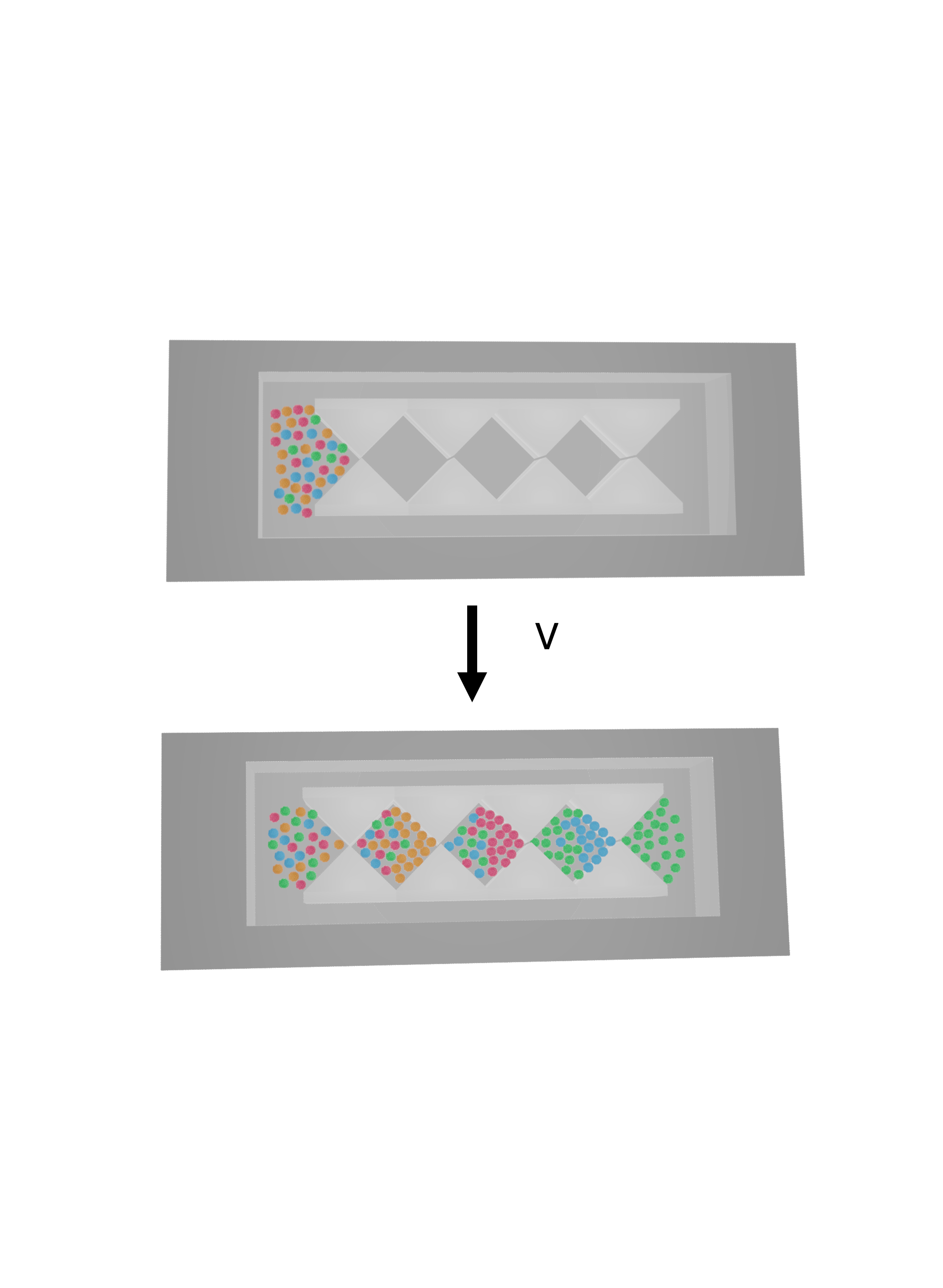

**Figure S1.** A truncated model of the microfluidic channel showing the separation of heterogeneous particles before (top) and after applying a voltage to separate particles (bottom).

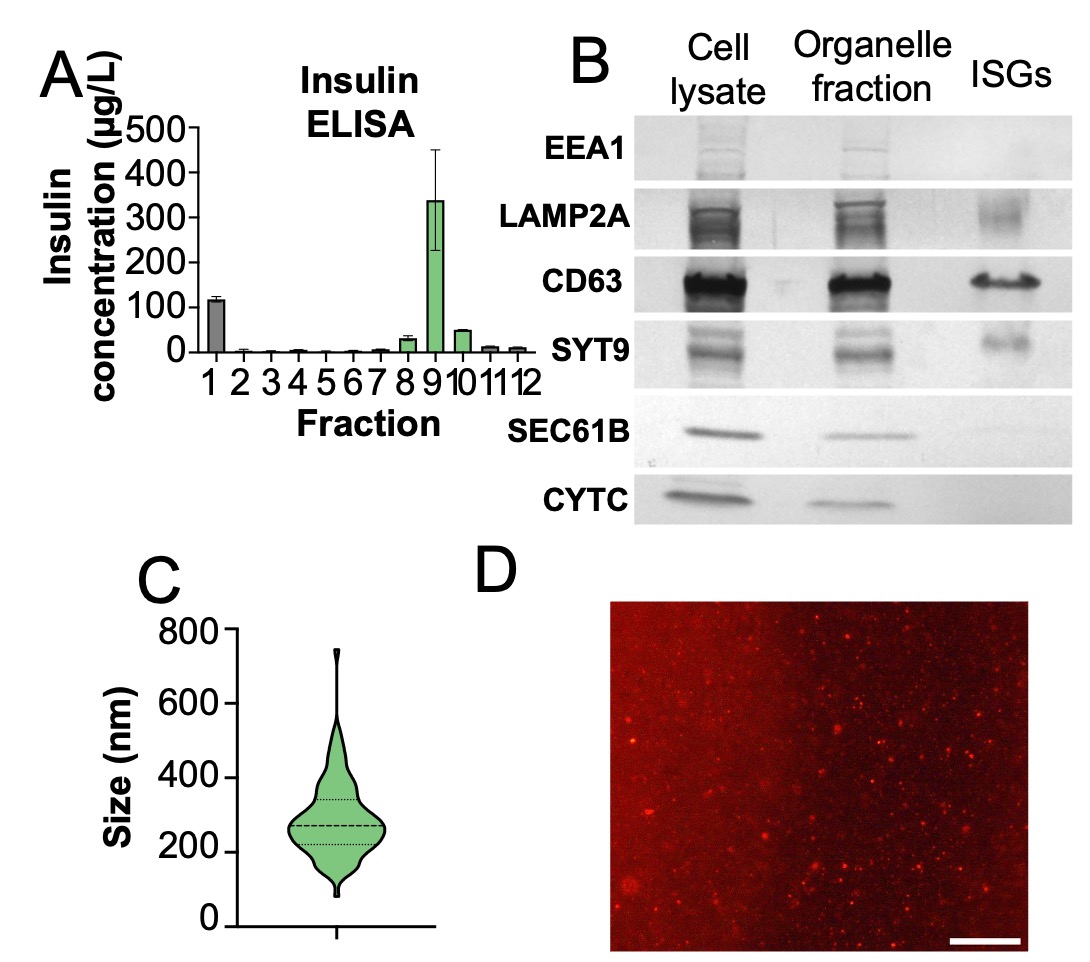

**Figure S2.** Validation of ISG isolation. **A.** Selection of fractions for use in separation experiments. Values are mean ± SEM. **B.** WB of cell lysate, organelle fraction, and isolated ISGs for determination of ISG purity. **C.** Size distribution of ISGs isolated from INS-1E cells. **D.** Fluorescence image of isolated ISGs. Scale bar: 250 µm.

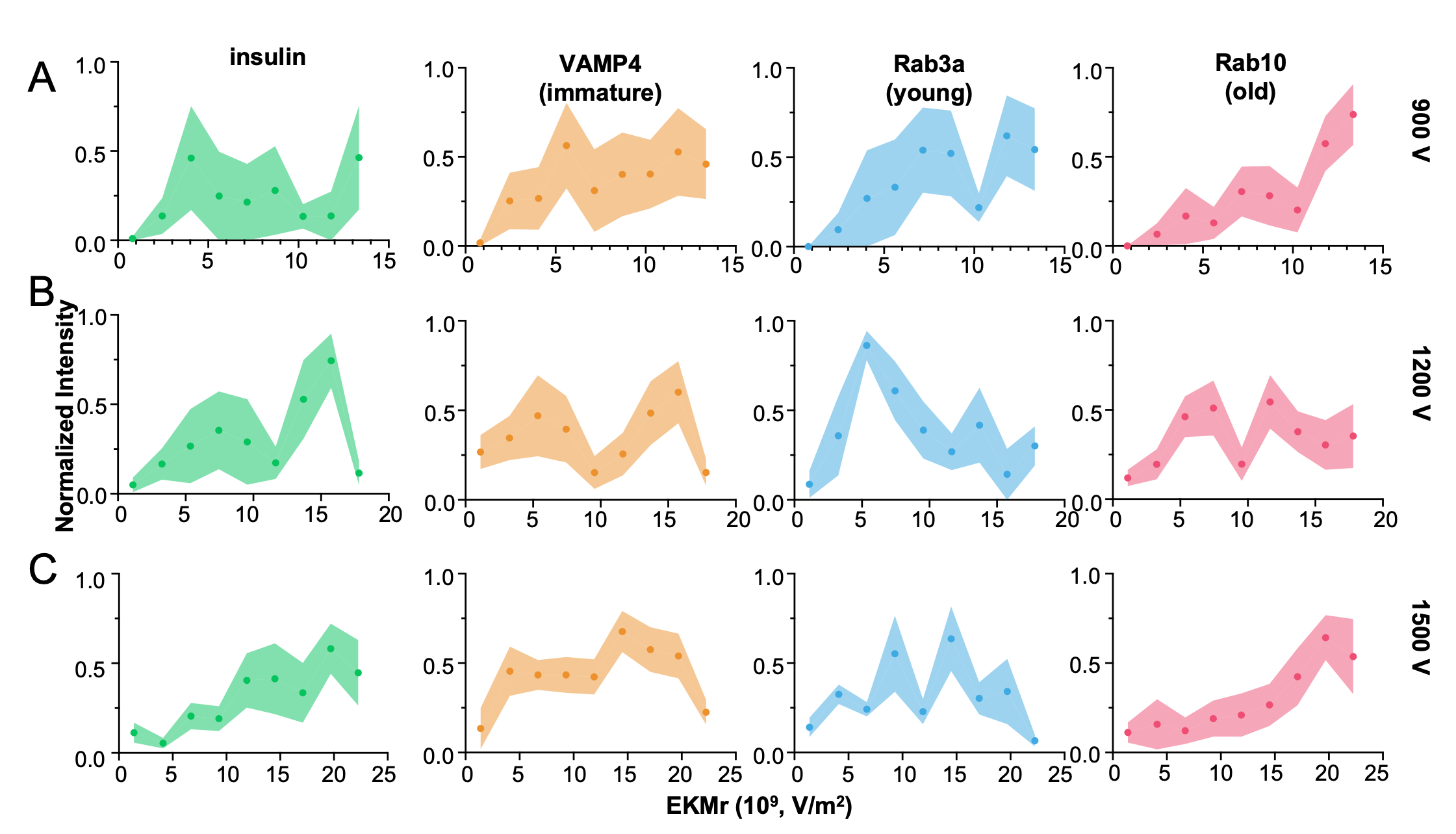

**Figure S3.** The EKMr distribution of different ISG subpopulations at **A.** 900 V (n=2-3 biologically independent experiments), **B.** 1200 V (n=3 biologically independent experiments), and **C.** 1500 V (n=3-4 biologically independent experiments). Values are mean ± SEM.

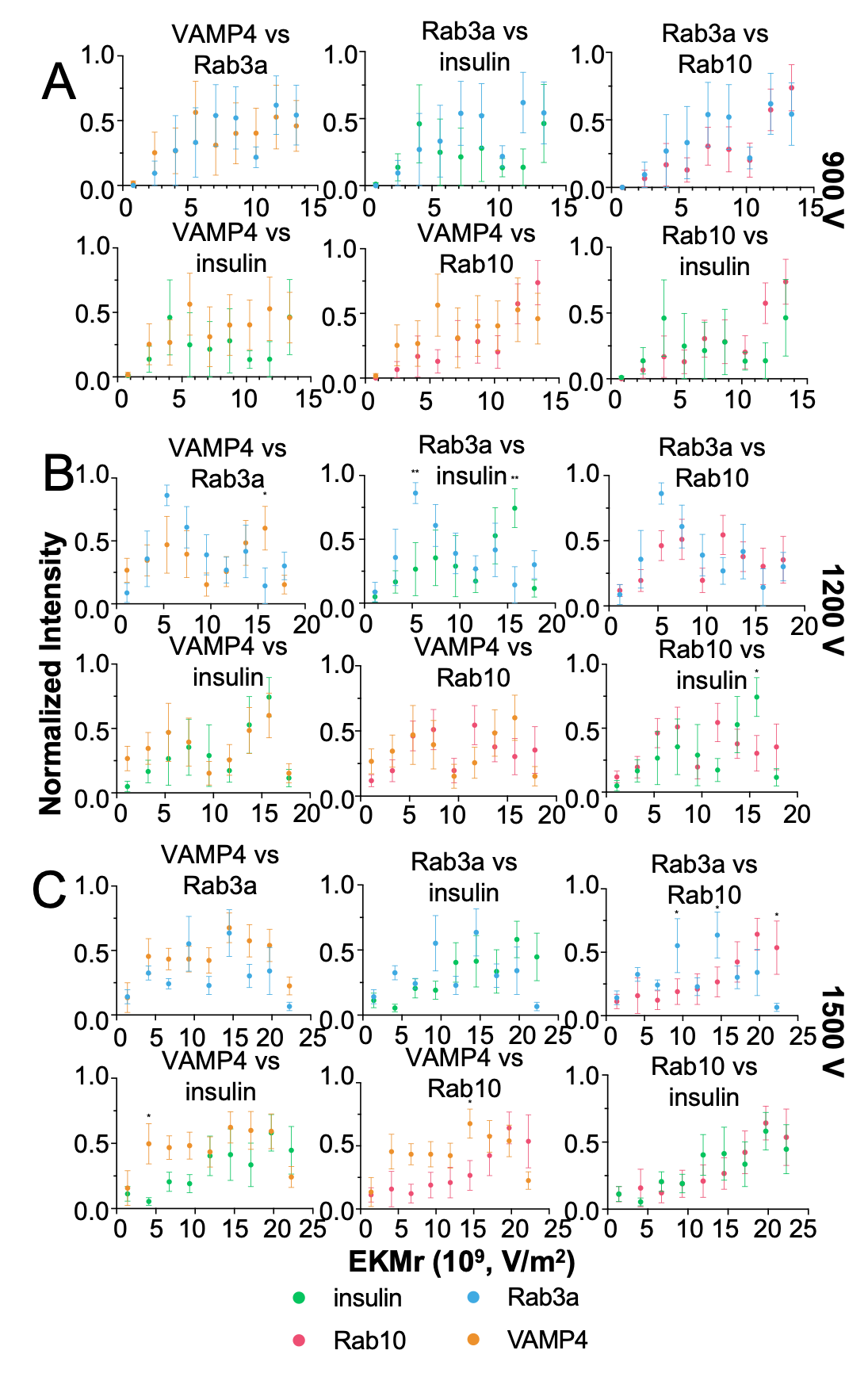

**Figure S4.** Comparisons of ISG subpopulation distributions at **A.** 900 V (n=2-3 biologically independent experiments), **B.** 1200 V (n=3 biologically independent experiments), and **C.** 1500 V (n=3-4 biologically independent experiments). Values are mean ± SEM (*p≤0.05, **p≤0.01 using ANOVA with Bonferroni post hoc multiple comparison correction).

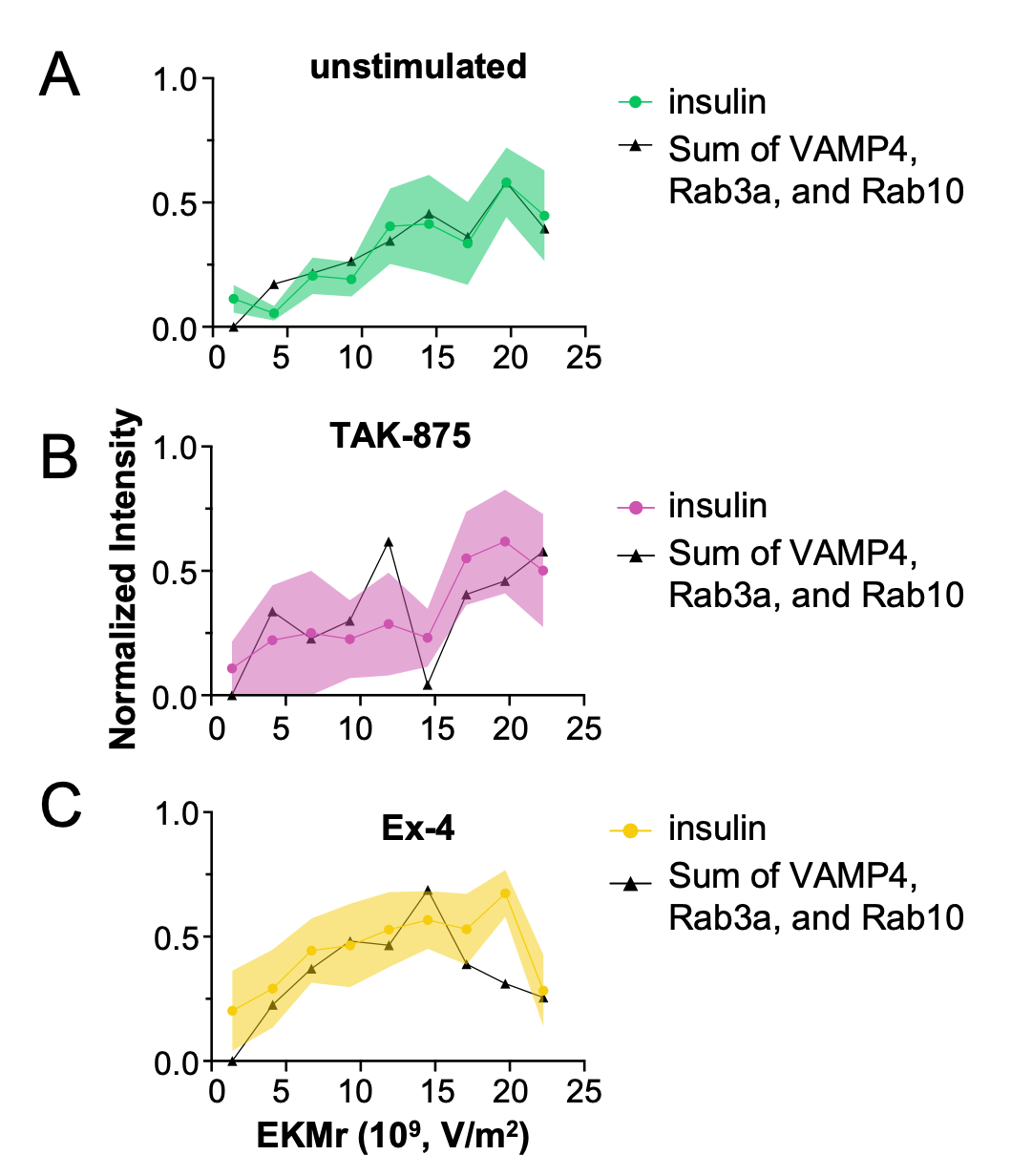

**Figure S5.** Plots of insulin are not significantly different from those of a linear combination of VAMP4, Rab3a, and Rab10. The average normalized intensity of each marker, excluding insulin, was summed and scaled to the same scale as the insulin plot from the same condition. These sums were plotted with insulin in the **A.** unstimulated, **B.** TAK-875-stimulated, and **C.** Ex-4-stimulated conditions. Values are mean ± SEM, and lines connect points for comprehensibility.

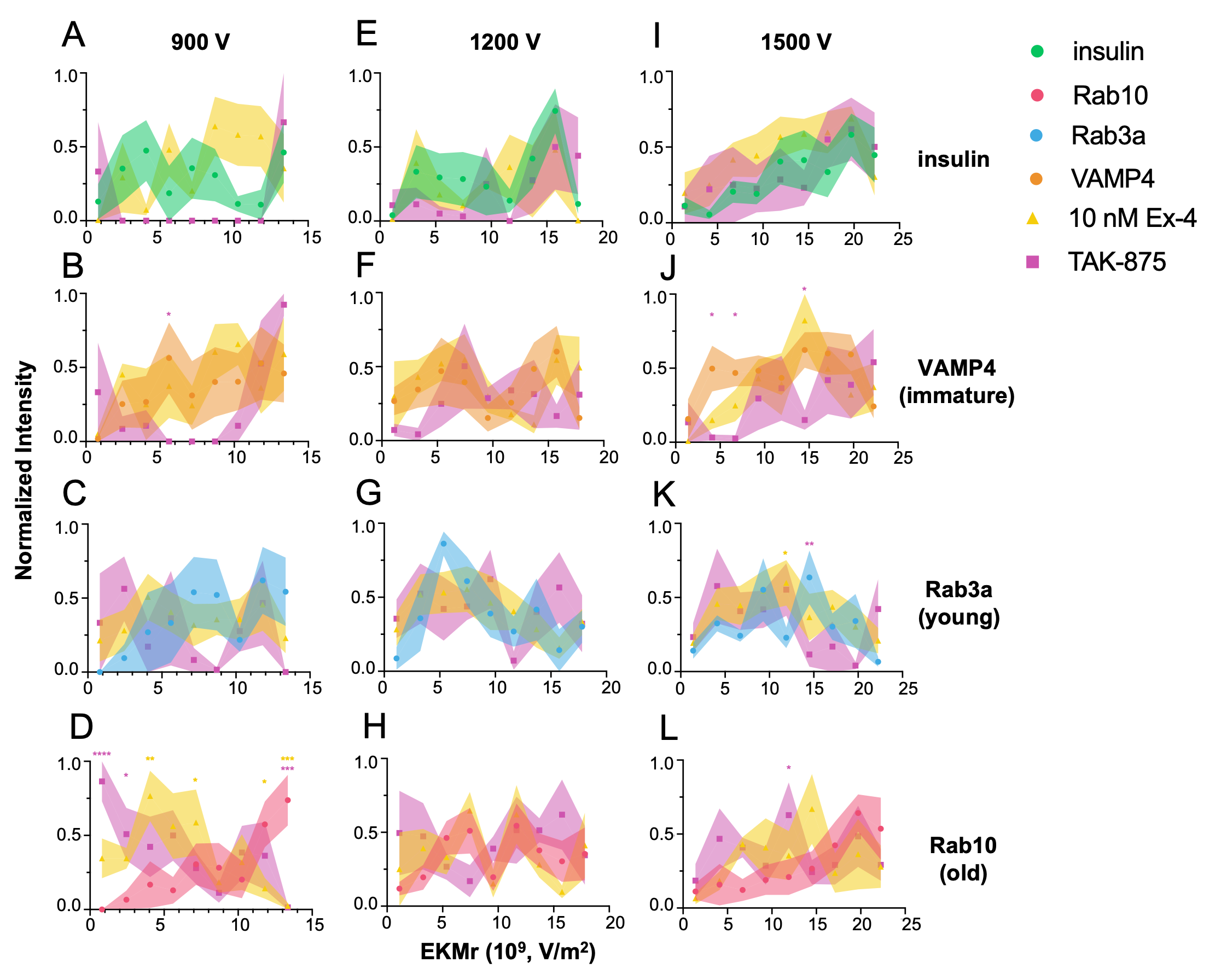

**Figure S6.** Changes in the EKMr distribution of each ISG subpopulation in response to TAK-875 and Ex-4 at **A-D.** 900 V (n=2-3 biologically independent experiments), **E-H.** 1200 V (n=2-3 biologically independent experiments), and **I-L.** 1500 V (n=2-4 biologically independent experiments). Values are mean ± SEM (*p≤0.05, **p≤0.01, ***p≤0.001 using ANOVA with Bonferroni post hoc multiple comparison correction. Comparisons made between stimulation and unstimulated conditions, symbols color-coded according to condition).

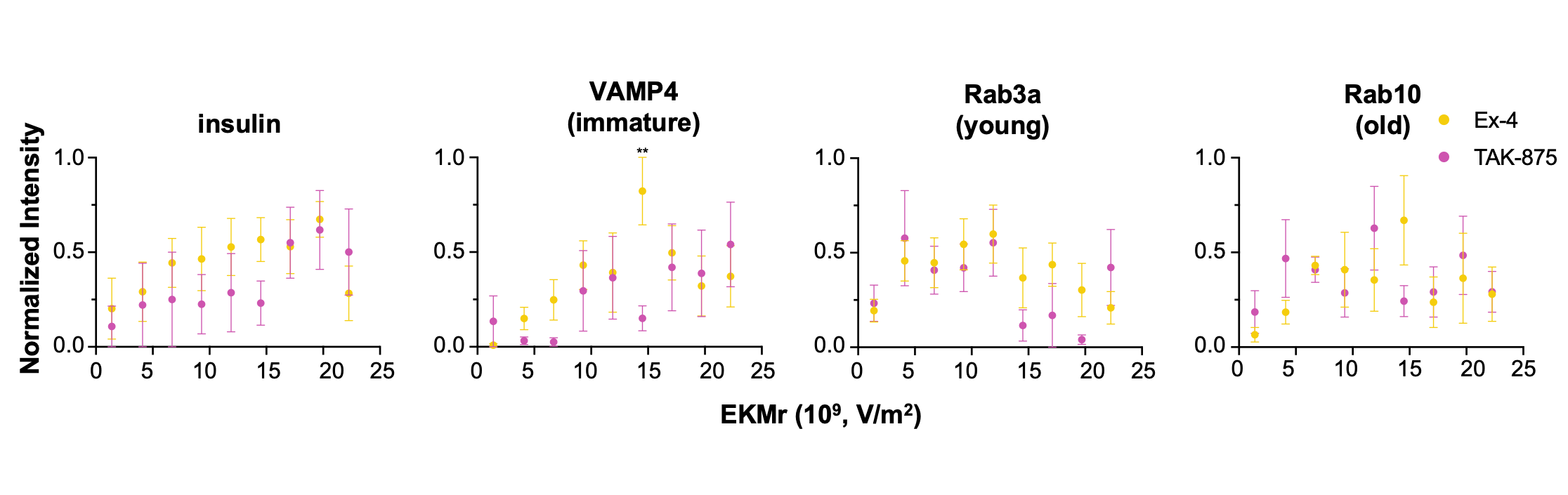

**Figure S7.** Differences in EKMr distributions between ISGs isolated from TAK-875 and Ex-4-stimulated cells at 1500 V (n=2-3 biologically relevant experiments). Values are mean ± SEM (*p≤0.05, **p≤0.01, ***p≤0.001 using ANOVA with Bonferroni post hoc multiple comparison correction).

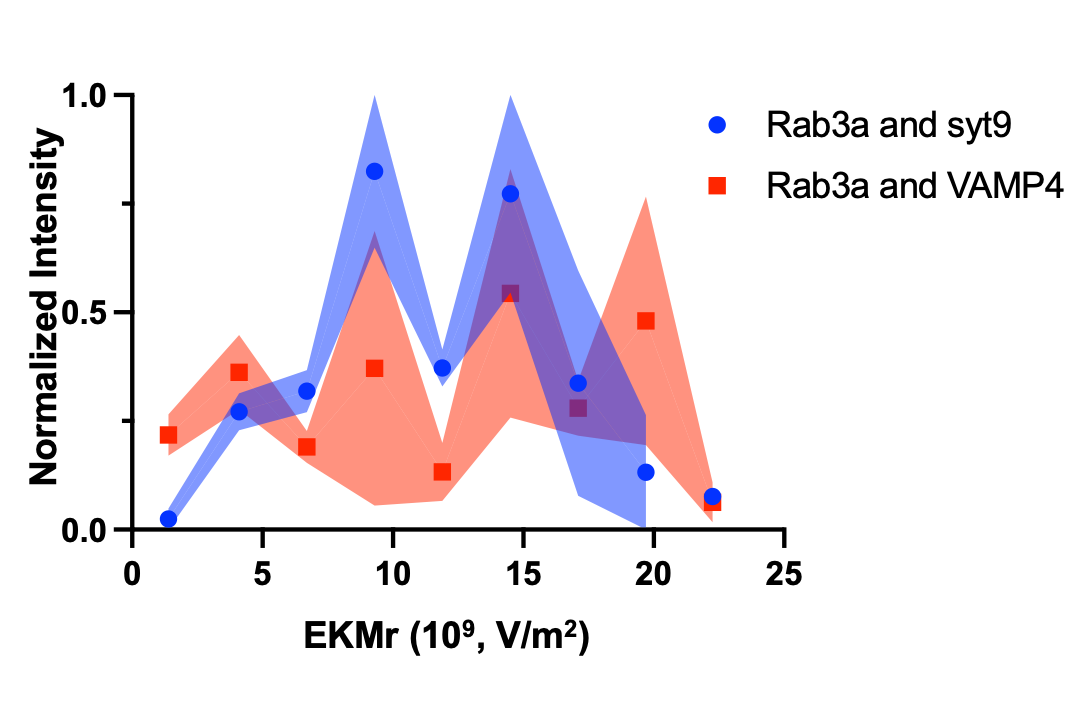

**Figure S8.** The EKMr distribution of young ISGs at 1500 V is not significantly affected by the differences in a second antibody label (syt9 or VAMP4, n=1-2 biologically independent experiments). Values are mean ± SEM.

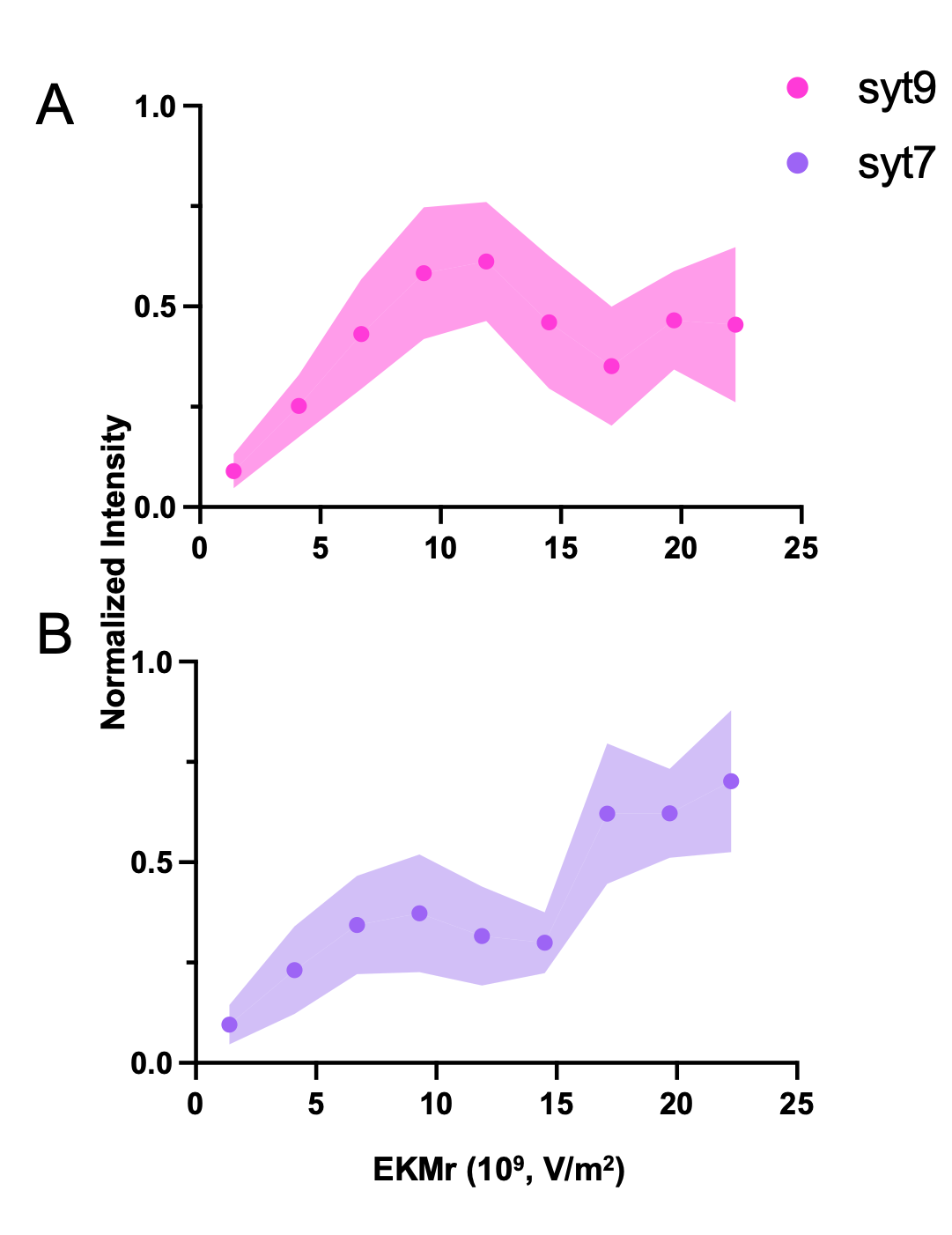

**Figure S9.** EKMr distributions of **A.** synaptotagmin 9 (syt9, n=5 biologically independent experiments) and **B.** synaptotagmin 7 (syt7, n=3 biologically independent experiments) at 1500 V. Values are mean ± SEM.

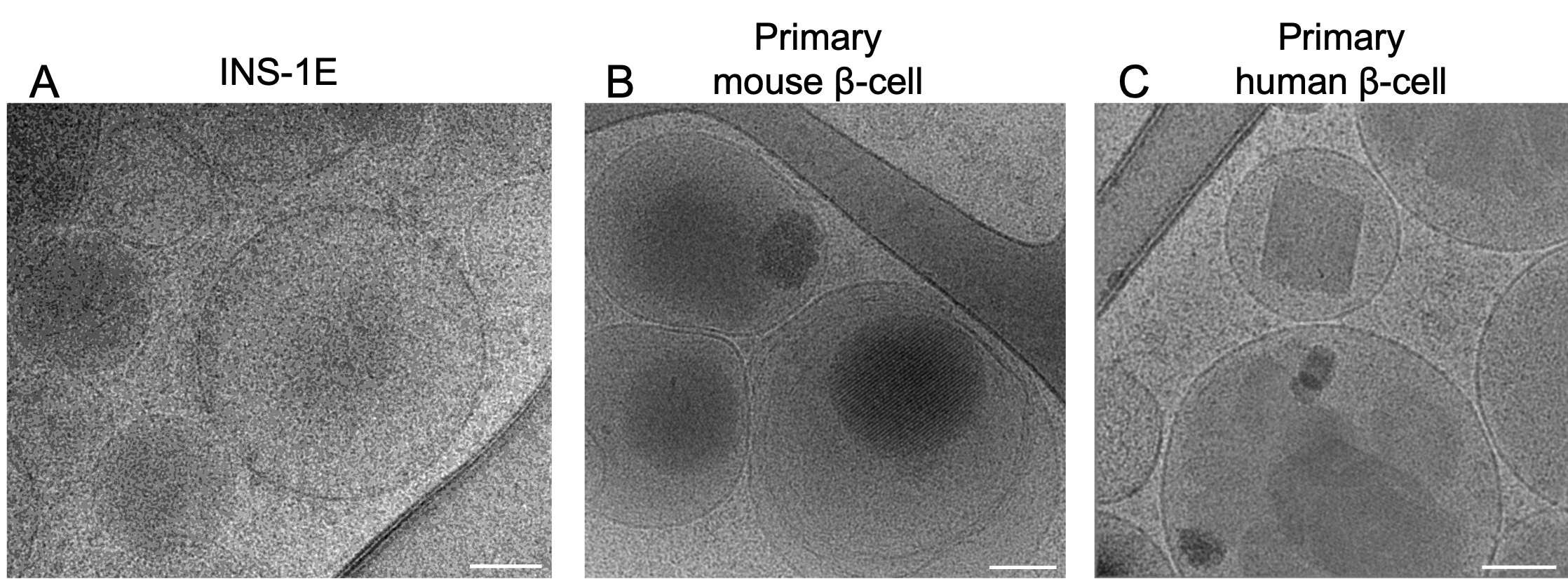

**Figure S10.** CryoET slices (**A-B**) and cryoEM images of ISGs inside INS-1E, primary mouse β-cells, and primary human β-cells. Magnification: **A-B.** 26,000x, **C.** 92,000x. Scale bar: 100 nm. A Gaussian blur of 1.0 nm was applied for clarity.

**Table S1.** Statistical comparisons between markers in the unstimulated condition.

**Table S2.** Statistical comparisons between the unstimulated condition and either the TAK-875 or Ex-4 condition.

**Table S3.** Statistical comparisons between the TAK-875 and Ex-4 conditions at 1500 V.
